# *Bifidobacterium breve* UCC2003 exopolysaccharide modulates the early life microbiota by acting as a dietary substrate

**DOI:** 10.1101/2019.12.17.879130

**Authors:** D Puengel, A Treveil, MJ Dalby, S Caim, IJ Colquhoun, C Booth, J Ketskemety, T Korcsmaros, D van Sinderen, MAE Lawson, LJ Hall

## Abstract

Members of the genus *Bifidobacterium* represent an important bacterial group for promoting health during early life. Previous studies have indicated that bifidobacterial exopolysaccharides (EPS) are involved in host interactions, with purified EPS also suggested to modulate microbe-microbe interactions by acting as a nutrient substrate. To further explore the role of EPS as a potential dietary component, we determined the longitudinal effects of bifidobacterial EPS on microbial communities and metabolite profiles using an infant model colon system. *Bifidobacterium breve* UCC2003 was utilised as a representative early life bifidobacterial strain, and a corresponding isogenic EPS-deletion mutant (*B. breve* UCC2003 EPS-). Initial transcriptomics analysis of the EPS mutant vs. parent *B. breve* UCC2003 strain highlighted differential expression in a discrete number of genes, including the *eps* biosynthetic cluster, though overall growth dynamics between the two strains were unaffected. Model colon vessels were inoculated with *B. breve* strains and microbiome dynamics were monitored using metataxonomic (via 16S rRNA sequencing) and metabolomic (via ^1^H NMR) approaches. Baseline early life microbiota profiles were similar between vessels, with persistence of *B. breve* (EPS+ and EPS-) observed between 0-36h. Within the EPS-positive vessel there was a significant shift in microbiome and metabolite profiles until the end of the study (405h); we observed increases of *Escherichia* and *Tyzzerella*, and short-chain fatty acids including acetate, propionate and formate, including further correlations between taxa and metabolites which were not observed in the EPS-negative vessel. These data indicate that the *B. breve* UCC2003 EPS is potentially being metabolised by members of the infant microbial community, leading to differential microbial metabolism and altered metabolite by-products. Overall, these findings may allow for development of EPS-specific strategies to beneficially alter the early life microbiota to promote infant health.

## Introduction

Members from the genus *Bifidobacterium* represent one of the dominant bacterial groups in the early life gut microbiota, with high levels associated with improved infant health [1-5]. Unlike the adult gut microbiome, the infant microbiome is less stable, and dietary change is proposed to lead to severe shifts in the abundance of major bacterial taxa over time [4, 6, 7]. The gut microbiota of breast-fed infants is dominated by *Bifidobacterium* (approx. 80% of the total community), which represents an important microbial pioneer or founder genus [8, 9]. In contrast, formula-fed infants have a more diverse microbiota and bifidobacteria comprise a smaller proportion (approx. 5-30%) [10]. The introduction of solid food at weaning marks a transition into a more complex microbiome, with a concurrent reduction in *Bifidobacterium* levels, likely due to the loss of milk as a sole dietary source. Notably, during these phases of significant dietary change, there is a shift in bifidobacterial species and strains, which may link to the wider repertoire of enzymes capable of digesting a more ‘adult’ diet [3, 5, 6, 11, 12].

Currently, only a small number of studies have explored the mechanisms by which *Bifidobacterium* species modulate the wider microbiota and/or provide benefits to the host. Several groups have provided data indicating that many *Bifidobacterium* species and strains produce exopolysaccharides (EPSs), which appear to have a variety of roles in microbe-host and microbe-microbe interactions [13]. Bacterial EPS are polymerised mono- or oligo-saccharides which form a diverse range of homo- or hetero-polysaccharides, that can be linked to the bacterial cell wall or secreted. Analyses of several *Bifidobacterium* species has revealed the presence of EPS gene clusters [14], with chemical analysis indicating glucose and galactose as major components, with very low levels of rhamnose also present throughout these structures [15, 16]. To date, most studies have focused on EPS-host interactions including; adhesion to host cells, and modulation of epithelial and immune responses [17-20].

Microbe-microbe interactions are a key feature of community structuring and are often mediated through cross-feeding of microbial-derived metabolites [21]. Interestingly, bifidobacterial EPS has previously been reported to act as a nutrient source for other bacteria within microbial communities. In two independent studies, Salazar and colleagues found that isolated EPS from human-associated *Bifidobacterium* species may act as a fermentable substrate for other members of the microbiota in faecal batch cultures (which was species and strain dependent) [22, 23]. More recently, this inter-microbial cross-feeding was also shown *in vitro*, with EPS from *Bifidobacterium animalis* subsp. *lactis* and *Bifidobacterium longum* promoting growth of *Bacteroides fragilis*, which correlated with increased short-chain fatty acid (SCFA) production [24].

Collectively, these studies suggest a potential role for EPS in microbe-microbe interactions, by acting as a substrate within gut microbiota cross-feeding networks. However, the exact role of bifidobacterial EPS in community re-structuring is currently unclear as previous studies supplemented vessels with purified EPS, rather than administrating an EPS-producing *Bifidobacterium* strain (which would mimic the actual gut environment). Here we explored the effects of *B. breve* UCC2003 (EPS-positive) and *B. breve* UCC2003del (EPS-negative) on the microbiota using an infant model colon system. We established differential gene expression and growth characteristics *in vitro* of each strain, prior to supplementation, and then performed longitudinal microbiota (via 16S rRNA) and metabolomic (via NMR) profiling, which indicated that EPS produced by *B. breve* UCC2003 can function as a potential dietary substrate that induces remodelling of the early life microbiota.

## Results

### Surface-associated EPS influences *B. breve* gene expression

Pure bacterial cultures were subjected to transmission electron microscopy to visualise the presence and absence of EPS prior to model colon experiments. Images indicated EPS-positive *B. breve* UCC2003 bacteria had a thicker and differentially stained cell wall (as indicated by arrows, **Fig 1a**) in contrast to the EPS-negative strain (i.e. *B. breve* UCC2003del); which is in line with previously published data [17].

**Figure 1:**
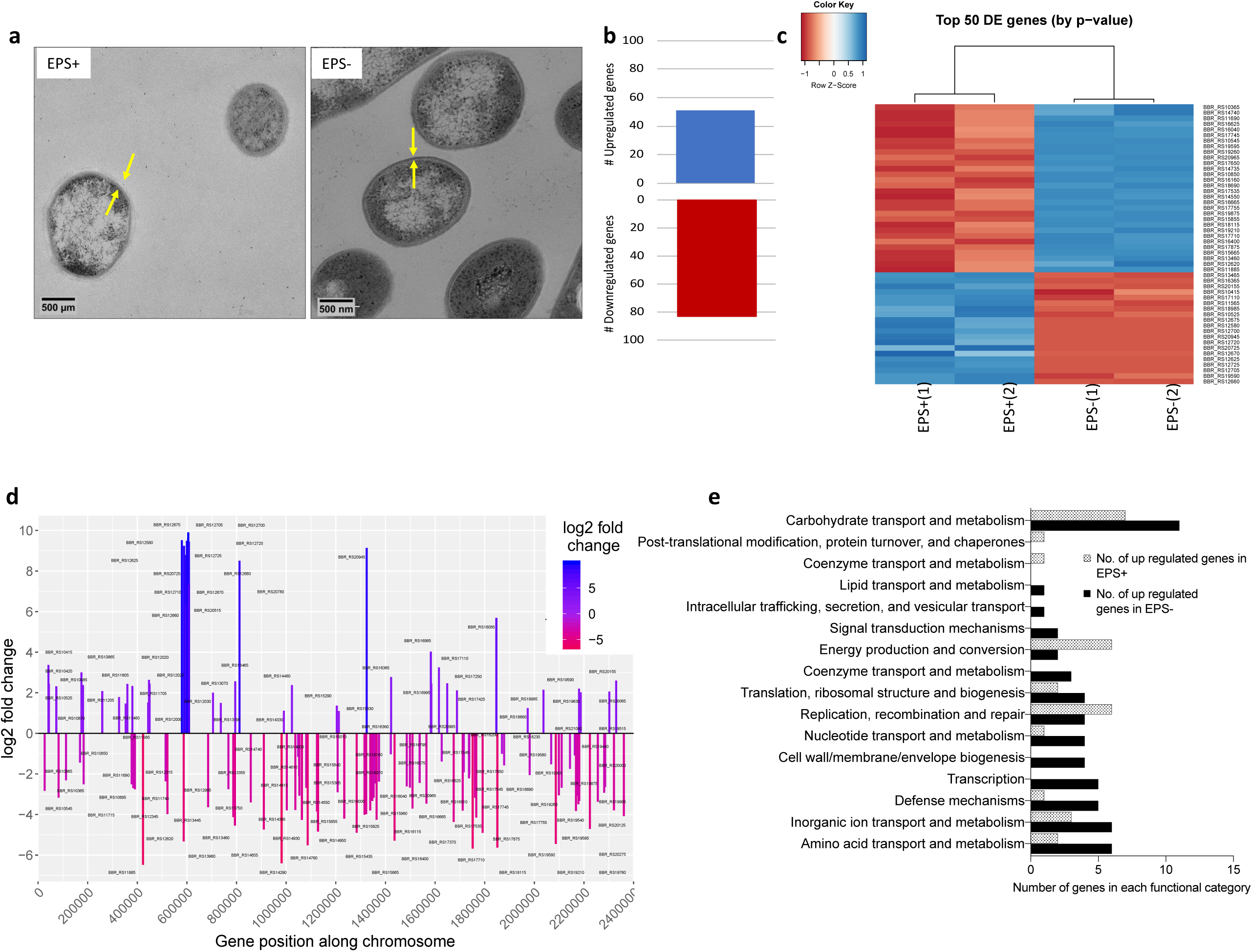
Characterisation of EPS-mediated *B. breve* modulation. **(a)** TEM of *B. breve* UCC2003 (EPS+) and *B. breve* UCC2003del (EPS-). Arrows note the EPS layer. **(b)** Total number of differentially expressed genes when comparing EPS+ to EPS- conditions, (two independent experimental repeats). **(c)** Top 50 differently expressed genes with an absolute log2 fold change ≥1 and p adj value ≤ 0.05. Hierarchical clustering of samples. Significance based on adjusted p value. Z scores are based on normalised, regularised log transformed and Z score transformed gene counts. **(d)** Bar plot of EPS+ vs EPS- differential gene expression (absolute log2 fold change ≥1 and p adj value ≤ 0.05) across the genome with gene name labels. **(e)** Number of differentially expressed genes in each functional category (EPS+ vs EPS-) according to EggNOG mapper annotation.

Previous work has suggested that bacterial EPS may act as a signaling molecule, and therefore its absence may alter bacterial gene expression [17]. To examine this, we grew *B. breve* EPS-positive and *B. breve* EPS-negative in culture until mid-exponential phase and isolated mRNA for transcriptome analysis. Principal component analysis (PCA) showed separation between EPS-positive and EPS-negative conditions (45% of variance, **Fig S1a**), in addition we found 51 upregulated and 83 downregulated genes (comparing EPS-positive to EPS-negative with an absolute log2 fold change ≥1 and p adjusted value ≤ 0.05; **Fig 1b** and **Fig 1c**). Annotation of regulated genes indicated high differential expression around the *B. breve* EPS cluster (**Fig 1d;** cluster at position ∼600,000). The EPS-related cluster was the only full gene cluster differently regulated between the two isolates during growth in rich media, suggesting that these strains are highly similar despite the lack of EPS structure on the mutant strain. Using EggNog mapper to functionally classify differentially expressed genes (from EPS+ vs EPS-cultures), the majority of genes were assigned to unknown function (18 genes up-regulated in EPS+ and 25 genes upregulated in EPS-, data not shown). In general, we found that the EPS-strain had a greater number of significant genes upregulated in metabolic pathways (including carbohydrate, nucleotide, inorganic ion, lipid and amino acid transport/metabolism) and bacterial cell growth (cell wall/membrane/envelope biogenesis and transcription; **Fig 1e**). Overall the EPS+ strain had fewer significantly differentially expressed genes, but in contrast to EPS-, EPS+ bacteria had a higher proportion of gene expression related to replication/recombination and repair, energy production and post-translation modification/protein turnover/chaperones and coenzyme transport and metabolism (**Fig 1e**). Collectively, this data suggests that the absence of EPS alters gene expression of many metabolic pathways, but importantly it does not influence overall growth kinetics (**Fig S1b**).

### *B. breve* EPS influences infant microbiome composition over time

To determine if bifidobacterial EPS acts as an additional nutrient substance, impacting cross-feeding activities, *B. breve* strains were inoculated into a complex infant model colon system (as depicted in **Fig 2a**). We used metataxanomic profiling, via 16S rRNA gene analysis, to probe kinetics of bacterial community changes induced by the presence of *B. breve* EPS-positive or EPS-negative strains over 405 hours of culture.

**Figure 2:**
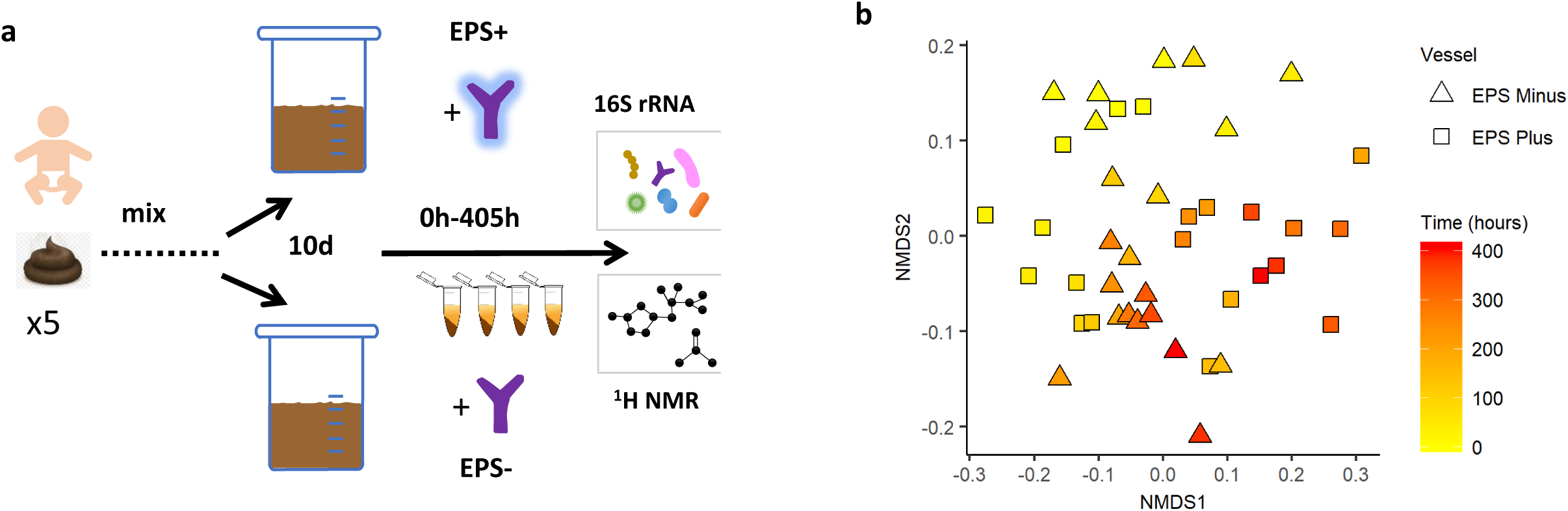
Experimental set-up and NMDS plots of EPS+ and EPS- vessels: **(a)** Five different stool samples from four different infants, age 6-12 months, were combined, processed and added to each Vessel (see Methods). Vessels were allowed to acclimatise for ten days, followed by inoculation with, Vessel A: *B. breve* UCC2003 EPS+, and Vessel B: *B. breve* UCC2003del EPS-. Aseptic sampling was performed from 0- 408h after inoculation and processed for 16S rRNA analyses and NMR. Samples were taken before inoculation (t=0), at time points (hours after inoculation) 6, 12, 24, 36 and from 48h to 408h every 24h. **(b)** NMDS plot using a Bray-Curtis dissimilarity calculation. For both vessels and separately (Fig S2). Changes over time are coloured from yellow (time point 0h) to red (time point 408h).

To offset the variability of microbial community dynamics that occur when stool samples from multiple infants are mixed and grown together in batch culture, all vessels were acclimatised for 10 days to stabilise ecosystems prior to *B. breve* supplementation at t=0 (**Fig S2**). *B. breve* levels were monitored over time by analysis of species-specific 16S rRNA abundance [25]; this method indicated that *B. breve* reads were only detectable at time 0h (following inoculation of each vessel) (**Fig S4a** and **S4b**). Sampling at 6h-36h post-supplementation showed presence of *B. breve* suggesting short-term colonisation within the infant model colon system. After a full media exchange (approx. 12h) *B. breve* accounted for approximately half of the total *Bifidobacterium* relative abundance within the EPS-positive vessel, and one-third of the total bifidobacterial abundance in the EPS-negative vessel (**Fig S4a** and **S4b**). This is perhaps unsurprising, since it has been previously reported that once stability has been reached in the microbiota, only one-third of individuals are colonized with *Bifidobacterium* after supplementation [26]. Despite short-term colonisation, overtime microbial diversity in each vessel shifted as indicated by NMDS plots of total taxonomic profiles (**Fig 2b** and **Figure S3a and S3b**), with specific changes in microbial composition and abundancy observed in each vessel throughout the experimental period (**Fig 3 & S5, Table S2 & S3**). Comparing the wider bacterial genus community profiles of the vessels indicated that both EPS-positive and EPS-negative conditions had similar core microbiomes, therefore we chose to focus our analysis on the top five genera in each condition. All vessels were dominated with a high proportion of (1) *Clostridium*, (2) *Bacteroides* and (3) *Erysipelatoclostridium* across all time points (**Fig 3a** and **3b**). Within the EPS+ vessel *Faecalibacterium* was the fourth most abundance genera across all time points; whilst bacteria belonging to the genera *Escherichia* were the fifth most prevalent at 0-36h, from 48-129h *Tyzzerella* was fifth, and *Enterococcus* from 144-408h (**Figure 3a and Table S2**). These shift in the top five genera within vessels corresponded to three unique microbial ‘phases’ (I, II, and III). Notably, presence of EPS (i.e. EPS+ vessel) correlated with different ecosystem structuring (**Figure 3b** and **Table S3**). Earlier time-points (0-36h) showed similar relative genus abundances when compared to the EPS-vessel; *Clostridium, Bacteroides, Erysipelatoclostridium, Faecalibacterium* and *Escherichia*. However, in the second phase from 48-120h, *Faecalibacterium* relative abundance was increased, with proportions of *Clostridioides* decreased (starting just before the second phase at 36h). Between 96h-120h, *Lachnoclostridium* and *Tyzzerella* had decreased read counts, whereas *Escherichia* increased. In the last phase, 144h-408h, *Clostridium, Bacteroides* and *Erysipelatoclostridium, Faecalibacterium* and *Tyzzerella* or *Escherichia* were most abundant, with *Enterococcus* relative abundance decreased, whereas the number of reads from *Tyzzerella* increased towards the end. These profiles indicate distinct and major changes in bacterial genera in response to *B. breve* UCC2003-associated EPS.

**Figure 3:**
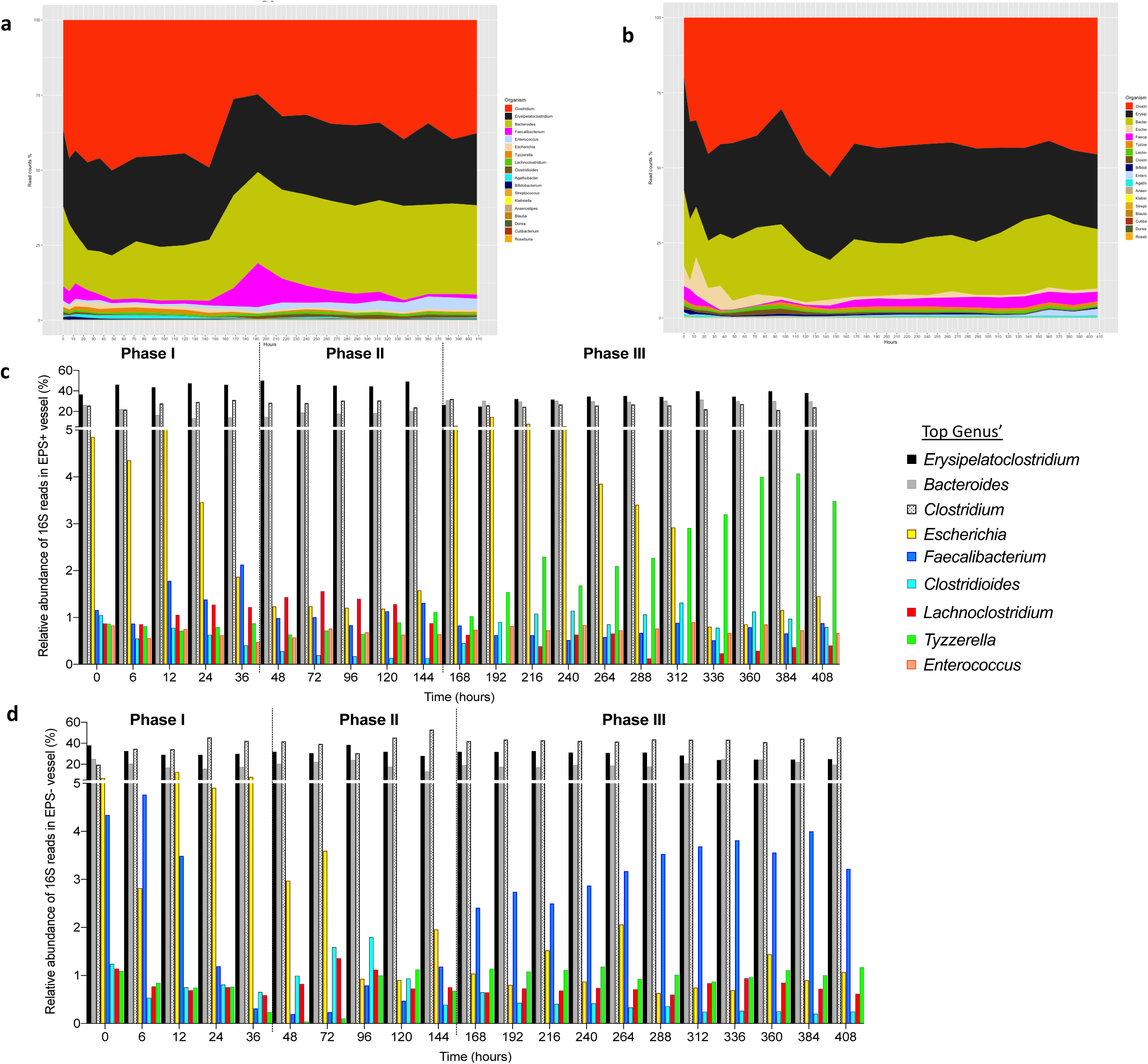
Microbiota profiling in EPS+ and EPS- vessels. Area plot of proportional read counts and total genus abundance of 16S rRNA gene analyses of **(a)** EPS+ and **(b)** EPS- model colon vessels. **(c)** EPS+ and **(d)** EPS- vessel changes (by phase) in the abundant genera.

Using Kendall rank correlation coefficient analysis, we next assessed if the presence of total *Bifidobacterium* correlated with other bacterial genera (**Fig S6**). Overall, we noted that most genera were found to be (weakly) negatively correlated in the EPS+ vs. the EPS– vessel, with a strongly positive co-occurrence of *Bifidobacterium* with *Faecalibacterium* in the EPS-vessel (p= 0.033; **Fig S6**). To determine putative EPS cross-feeding partners we also analysed for co-occurrences with *B. breve* (i.e. specific for EPS+/EPS-supplemented strains) we found that, although not significant, there was a trend for more positive correlations with several genus in the EPS+ vessel including *Streptococcus* and *Ruminococcus.* In general, we observed fewer positive correlations EPS-*B. breve* correlations, with the genus *Dorea* significantly negatively correlated (**Fig S6**). Collectively, this might suggest that the presence of EPS is responsible for driving changes in microbiome profiles, likely due to the fact it can function as a potential metabolite for other members in the microbiota.

### Metabolite profiles shift in response to bifidobacterial EPS

Using ^1^H NMR, we next sought to examine if the taxonomic compositional changes observed were also linked to overall metabolic changes. In agreement with the microbiota profiles, both vessels had similar starting metabolite concentrations prior to the introduction of *B. breve* (**Fig S7a**). The most abundant metabolites in each vessel were formate, succinate, ethanol, and bacterial fermentation by-product SCFAs: acetate, butyrate and propionate (**Fig. 4a-d; Table S4**). Proportions of all SCFAs increased over time independent of the presence/absence of EPS (**Fig 4c-d**) [27]. We also observed that within the EPS-vessel, the concentration of ethanol increased as time progressed, whilst the EPS+ vessel appeared to have decreasing levels. The abundance of the simple organic compound formate increased in abundance until 120h within the EPS+ vessel, then dropped in the third phase (144-406h) but remained stable in the EPS-vessel. We observed that the sugar compound succinate decreased with time in the EPS-vessel, whilst levels were unchanged in the EPS+ system. Taken together these results further suggest that addition of bacteria with an EPS structure alters bacterial function and metabolic output.

**Figure 4:**
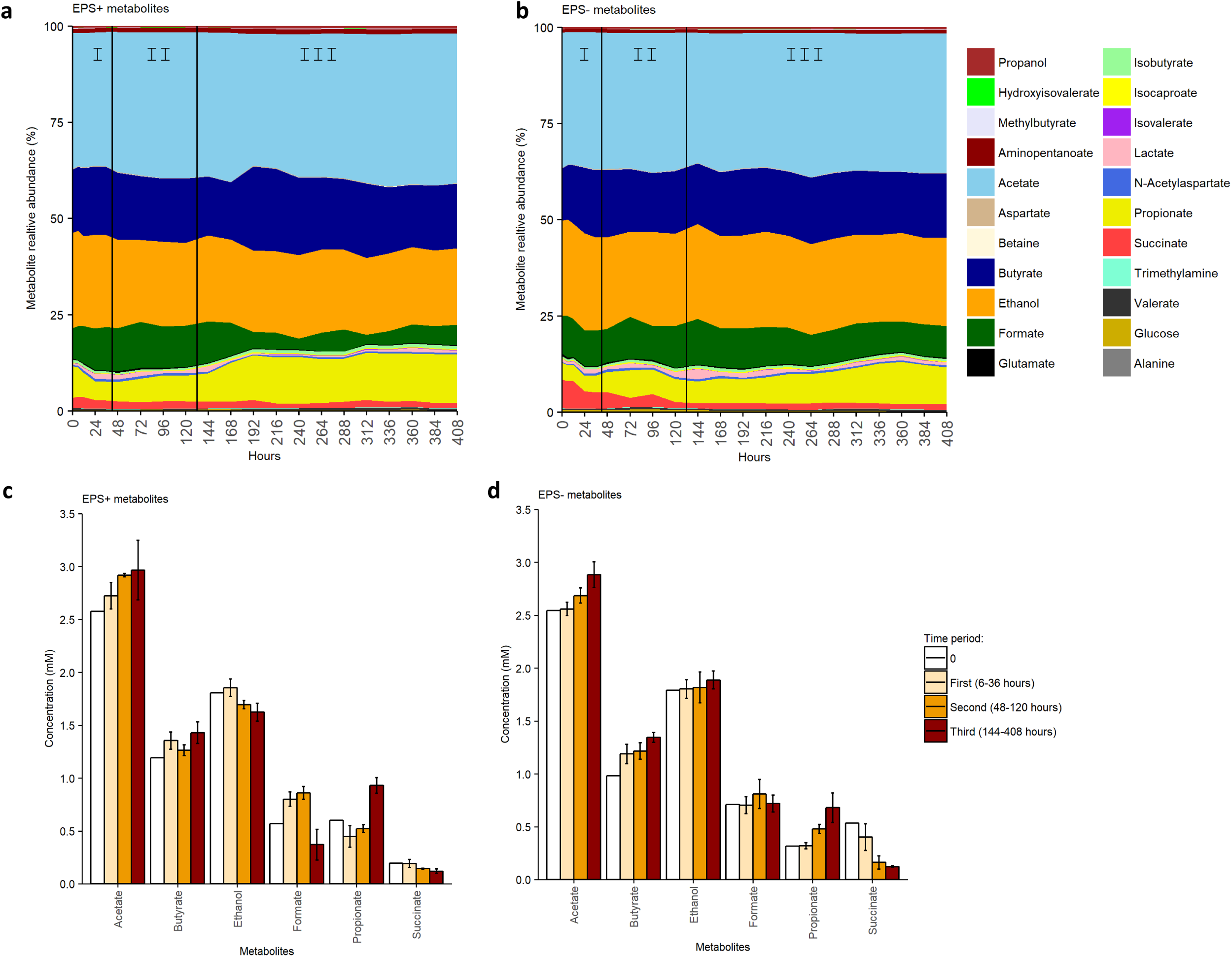
Metabolites profiling of EPS+ and EPS- vessels over time. Metabolite proportions as measured via 1H NMR, detected in **(a)** EPS+ and **(b)** EPS- vessels of model colon experiment calculated to 100%. Individual growth phases are denoted on each graph. The mean concentration of the most abundant (nM) metabolites within each time period (at 0h, 6-36h, 48-120h and 120-408h) for each vessel given either **(c)** *B. breve* UCC2003 EPS+ and **(d)** *B. breve* UCC2003del EPS-. Bars represent mean concentration for each metabolite over the specified time period, error bars represent standard deviation.

Direct comparison of metabolic and microbiota analysis using the Spearman correlation analysis identified changes in bacterial profiles that could be linked to EPS-mediated metabolic changes (**Fig 2a, 5 & S7b**). Overall, we noted more significant associations in the EPS+ vessel, and therefore chose to focus on the most abundant metabolites with the top 10 bacterial genera across the experimental period. The strongest positive correlations (i.e. both genus and metabolite increasing; p≤0.001) were observed in the EPS+ vessel between *Bifidobacterium* and ethanol; *Tyzzerella, Escherichia, Clostridium, Agathobacter* and formate; *Enterococcus, Bacteroide*s and propionate and, *Escherichia* and succinate (**Fig 5; Table S5**). The presence of EPS+ *B. breve* was positively associated the SCFA butyrate and *Clostridioides* (and negatively associated in the EPS-vessel). Additionally, propionate was significantly negatively correlated with the proportion of *Tyzzerella* and Agathobacter in the EPS+ vessel and *Clostridioides* in the EPS-vessel. The apparent differences in the metabolome with bacterial genus correlations indicate distinct and related changes due to the presence (or absence) of EPS; and may possibly infer active EPS metabolism (when present) by members of the wider early life microbiota.

**Figure 5:**
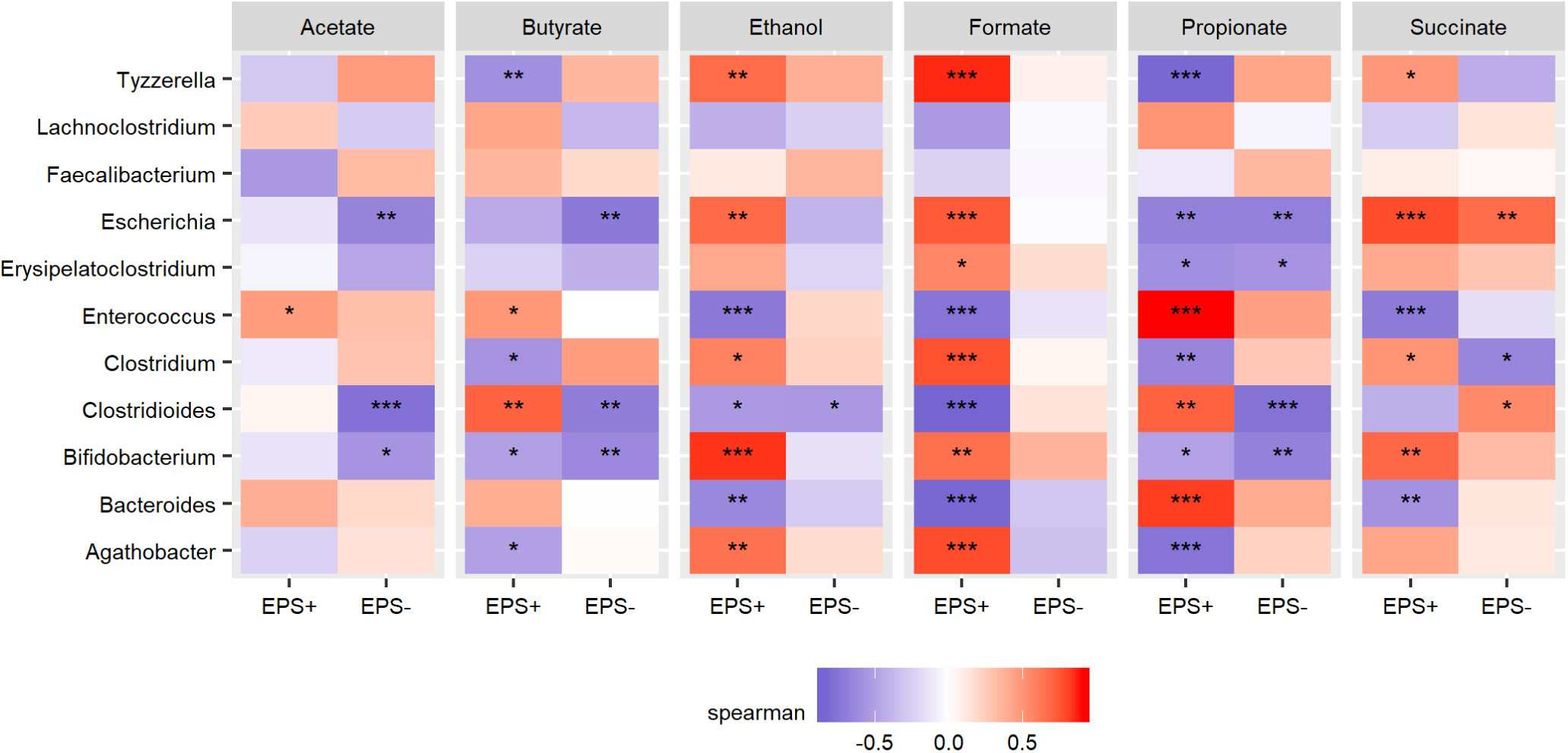
Spearman correlation between the top 10 bacterial genus and top six abundant metabolite in each vessel (EPS+ vs. EPS-). *p < 0.05; **p<0.01; ***p<0.001. The P-values are adjusted for multiple comparison using Benjamini & Hochberg by genus, and by each vessel (EPS+ and EPS-).

## Discussion

Previous studies have indicated that bifidobacterial EPS plays a key role in mediating beneficial microbe-host interactions [1]. Earlier studies have also suggested that EPS potentially acts as a dietary component, facilitating cross-feeding within microbial communities [22]. EPS molecules are large surface-bound polymers surrounding the cell wall of many Gram positive bacteria, including *Bifidobacterium* species [13]. Expression of EPS may influence overall growth of *B. breve* due to increased energy expenditure, and/or EPS modulation of bacterial gene expression, which may impact profiles within complex microbial ecosystems [18]. RNASeq analysis indicated regulation of the EPS biosynthetic gene cluster (including glycosyltransferases) suggesting that EPS may act to modulate its own production, as indicated by TEM images [17]. Indeed, previous work in *Bacillus subtilis*, indicates that presence of EPS promotes the phosphorylation of a glycosyltransferase in the biosynthetic pathway, thereby stimulating the production of EPS [28]. We found that *in vitro*, presence of EPS did not significantly alter overall gene expression or growth, indicating any changes observed from the addition of *B. breve* UCC2003 EPS-positive/negative to a complex microbiota are due to presence (or absence) of EPS, rather than inherent cell response differences due to the additional exopolysaccharide motif.

Significant variations are often observed between infant microbiotas, particularly during weaning when ecosystems are undergoing rapid change in response to a changing nutritional environment (between 6-12 months of age). Metataxonomic analyses of the microbial ‘baseline’ model colon profiles revealed an expected weaning infant core microbiome; namely dominance of the *Bacteroides* genus, and members of the phylum *Firmicutes (e.g. Clostridium* and *Erysipelatoclostridium)*, with reduced abundance of *Bifidobacterium* [3, 5, 11]. This developmental window represents an important dietary transition period for infants; moving from milk (breast or formula) to solid food, therefore access to additional microbial-derived nutrients, such as EPS, may contribute to driving ecosystem re-structuring [4]. Indeed, our research indicates that *B. breve* UCC2003 can modulate the infant microbial community structure and metabolism, and in line with previous studies we noted alterations in microbially-derived SCFAs; acetate, propionate and butyrate, although we did not detect the previously reported EPS driven acetate to propionate ratio change [4, 19], which may be due to experimental and inoculating infant microbiota differences.

Previous studies on *B. breve* EPS has identified glucose and galactose as main components of EPS, which is similar to that proposed for UCC2003 EPS structure (includes glucose, galactose and/or the N-acetylated versions of these two sugars in different ratios or composition) [13, 15-17]. The large molecular weight and structure of these EPS may prevent host-associated digestion, allowing them to act as prebiotics within the colon (similar to inulin and fructooligosaccharide) [19, 29, 30]. Interestingly we found that *Escherichia* increased over time and was highly correlated (positively) with *Bifidobacterium* in the EPS+ vessel from 0-36h (when *B. breve* could be detected). *Escherichia* is known to encode β-galactosidase genes that may allow degradation of the sugars present in EPS polymers [31, 32]; potentially correlating with the increases in metabolites such as ethanol, propionate and formate also during this growth phase [33, 34]. The infant-gut associated bacteria *Tyzzerella* [3], was also dominant in the EPS+ vessel (at 48h) and may therefore use EPS as a key nutritional supplement. Although little is known regarding the metabolism of members the *Tyzzerella* genus, many other members of the family *Lachnospiraceae* possess glycoside hydrolases used for the metabolism and transport of oligosaccharides into the cell [35]. Furthermore, another member of the *Lachnospiraceae* family, *Ruminococcus*, degrades intestinal mucins (similar in structure to EPS) liberating metabolites for other members of the microbiota to consume [36]. Thus, the positive correlation of *Tyzzerella* with formate in the EPS+ vessels suggests that, similar to other members in the *Lachnospiraceae* family, *Tyzzerella* may be consuming EPS and producing formate as a fermentation end-product [37]. Moreover, we observed several other strong negative correlations between different genus (e.g. *Dorea*) and *B. breve* UCC2003, which also suggests antagonistic networks may be EPS-dependent, but further studies are required to understand the mechanisms promoting these inverse relationships.

As highlighted above, the infant microbiota represents a dynamic ecosystem, with complex feeding networks at play, which may also be acting in response to EPS metabolism. We noted several potential wider cross-feeding networks (rather than direct EPS digestion interactions) during different growth phases. Specifically, utilisation of *B. breve* EPS by primary degraders e.g. *Escherichia* producing N-acetylaspartate may also facilitate growth of secondary degraders such as weaning-associated *Faecalibacterium*, important butyrate producers, as these metabolites correlate respectively with the two genera [38-41]. Another possible interaction is via predicted production of formate by *Tyzzerella*, that may be metabolised by *Faecalibacterium*, as has been described in an *in silico* biofilm model [42] (**Fig S8)**. Delineating intricate and complex cross-feeding relationships is difficult and cannot be predicted from non-community cultures, as strains grow, and behave differently when grown in co-culture, compared to those grown individually [43]. Further studies, possible utilising isotope labelling, would be required for targeted identification of metabolites impacting these important EPS-associated cross-feeding networks [44].

To our knowledge, this research demonstrates for the first time, the influence of *B. breve* UCC2003 EPS on the infant microbiota using an *in vitro* model colon system. Although a dynamic system, with media constantly replenished and waste removed, we observed short-term colonisation that contributed to altered bacterial composition (i.e. abundance of *Escherichia* or *Tyzzerella*) and metabolite profiles (i.e. acetate, propionate, formate or butyrate) within the EPS-positive *B. breve* UCC2003 inoculated vessel. These combined data suggest EPS may act as a nutrient source for the early life microbiota thereby driving changes in microbial community structure. Our results, in combination with previously published work [22, 23] sugget that early life supplementation with EPS-positive *Bifidobacterium* strains may promote infant well-being, by aiding and supporting beneficial infant microbiota development. Finally, this work, including the identification of microbial associations and metabolites within a healthy microbiota, could be used as a platform for future nutritional- and microbial therapies development strategies.

## Materials and Methods

### Bacterial Strains and Growth Conditions

*B. breve* UCC2003 strains used in this experiment included *B. breve* UCC2003 (i.e. EPS-positive, and isolated from infant stool), and EPS deletion mutant *B. breve* UCC2003del (i.e. EPS-negative, [17]). All strains were cultured in reinforced Clostridium medium (RCM) and deMan-Rogosa-Sharpe (MRS) medium and supplemented with 0.05mg/mL L^−^cysteine (L-cys) HCl anaerobically at 37°C for upwards of 48h. When measuring growth kinetics samples were taken every 2 hours from 0h to 12h and 24h to 30h, with a measurement after 48h of growth. Optical density (600nm; OD_600_) and pH values were recorded at each time point. For colony forming units (CFU) per mL determination, samples were plated onto RCM agar plates containing 0.05mg/mL L-cys-HCl and grown anaerobically for 48h.

### *B. breve* transcriptomics and bioinformatics analysis

To identify potential EPS modulated genes, pre-cultures grown in MRS media overnight (as above) were used to inoculate fresh cultures and cells were harvested at exponential (approx. 8 h) phase, PBS washed, and RNA extraction was performed immediately. The cells were resuspended in 600 µL RLT lysis buffer containing 8 µL β-mercaptoethanol. The suspension was transferred to Fastprep Lysing Matrix E tube (MP Bio) and was homogenised using FastPrep-24 instrument (MP Bio) at 6.0 m/s for 3 × 1 min with 5 min resting intervals on ice. Samples were centrifuged at 14,000 × *g* for 10 min and RNA was extracted using Qiagen RNeasy mini plus kit (Qiagen) according to manufacturer’s instructions. The RNA quality and concentration were determined using Agilent 2100 Bioanalyzer (Agilent Inc.). Only samples with RIN values above 8 were sequenced. Isolated RNA was processed with Ribo-depletion, and samples sequenced on HiSeq V4 75bp using non^−^stranded, paired end reads.

For computational analysis, the quality of stranded reads was assessed by FastQC software (version 0.11.8) [45]. Reads were aligned against the full nucleotide sequence of *B. breve* UCC2003 (RefSeq: NC_020517). Alignment and quantification were performed using Kallisto (version 0.44.0), the quantified read data was normalised and differential expression analysis was conducted using DeSeq2 (version 1.22.2) [46, 47]. Genes with an absolute log2 fold change ≥1 and *p* adj value ≤ 0.05 were considered to be differentially expressed. PCA plots, heatplots and plots of differential expression across the genome were generated using custom R scripts. All genes in the *B. breve* UCC2003 genome (RefSeq: NC_020517) were functionally annotated with categories and descriptions using EggNog-mapper (version 4.5) [48]. Custom R scripts were used to test for enrichment of EggNog functional categories in the upregulated and downregulated differentially expressed genes. This analysis was carried out using gene set enrichment functions of the R package clusterProfiler (version 3.10.1) [49]. All raw reads were deposited in European Nucleotide Archive: PRJEB35291.

### Transmission electron microscopy (TEM)

Strains were grown as described above, and after 48h growth, cells were centrifuged at 4500 × *g* at 4°C for 10min with low break. Bacterial pellets were resuspended in 1mL of fixative comprising 2.5% glutaraldehyde in 0.05M sodium cacodylate pH 7.2. Fixation was carried out for 1.5h at room temperature after which, cell suspensions were centrifuged (7.5xg rpm, 3min) and washed in 0.05M Sodium Cacodylate buffer x3 (10min). After final centrifugation, cell pellets were mixed 1:1 with molten 2% low gelling temperature agarose, solidified by chilling, and then chopped into 1mm^3^ pieces. Sample pieces were post fixed in 1% OsO_4_ for 2h followed by washing x3 (15min) in deionised water. Sample pieces were then dehydrated through an ethanol series (30%, 50%, 70%, 90%, 100% x3) for at least 15min in each ethanol dilution. Samples were infiltrated with a 1:1 mix of LR White medium grade resin to 100% ethanol, followed by a 2:1 and a 3:1 mix and finally 100% resin, with 1 hour between each change. This was followed by two more changes into fresh 100% resin, with periods of 8 hours between. Four blocks/sample were put into Beem capsules (Size 00) with fresh resin and polymerised for 24h at 60°C. Sections approximately 90nm thick were cut using an ultramicrotome with a glass knife, collected on formvar/carbon coated 200 mesh copper grids, and stained sequentially with 2% uranyl acetate for 1h at room temperature; and 0.5% lead citrate-tribasic trihydrate for 1min at room temperature. Deionised water washes were performed (x5) following each of the staining steps. Sections were examined and imaged in a Talos F200C transmission electron microscope at 200kV with a “Gatan One View” digital Camera.

### Infant model colon system

Faeces were collected from healthy, full-term breast-fed infants (aged from 7-month to 1-year old) in accordance with protocols by the National Research Ethics Service (NRES) approved UEA/QIB Biorepository (Licence no: 11208) and the Quadram Institute Bioscience Ethics Committee. One-gram samples from five frozen stool samples were homogenized with 5mL reduced PBS (500µL of 3% L-cys HCl in 50mL PBS), filtered (70μm) and evenly distributed to each vessel. Model colon system media was made according to Cinquin et al., and included a vitamin solution (pantothenate 10mg/L, nicotinamide 5mg/L, thiamine 4mg/L, biotin 2mg/L, vitamin B12 0.5mg/L, menadione, 1mg/L and p-aminobenzoic acid 5mg/L) as described in Gibson and Wang [50, 51]. The continuous-fed batch cultivation was performed in 1.4L bioreactors (Multifors) and monitored with Eve® software (Infors AG). Parameters were set according to [50]; briefly, vessels were maintained at 37°C, pH 6.7 and anaerobic, with a media exchange occurring every 12h (retention time). All vessels were given 10 days to permit equilibrium of the microbiota to the *in vitro* vessel environment.

For inoculation of *B. breve*, strains were grown in the infant gut media. Overnight cultures were then centrifuged at 4500 × *g* for 10 min at 4°C, washed with PBS and then inoculated into each vessel. *B. breve* UCC2003 EPS-positive 4.4 × 10^8^ CFU/vessel, and *B. breve* UCC2003del EPS-negative 2.4 × 10^8^ CFU/vessel was added. Samples were snap frozen in liquid nitrogen at each time point t=0, 6, 12, 24, 36 hours post inoculation with *B. breve*, and once a day from that time point onward.

### DNA extraction, 16S rRNA library preparation, sequencing, and bioinformatics analysis

For genomic DNA extraction of the samples FastDNA® Spin Kit for Soil (MP Biomedical) was used accordingly to manufactures instructions with an additional two bead-beating runs of 60s and setting 6.0m/s. DNA quantification was determined with the Qubit® 2.0 Fluorometer using the broad range Qubit® Assay Kits for DNA, RNA and Protein following manual instructions, and DNA was normalised to a final concentration between 4–20ng/µL prior to 16S PCR analysis.

V1 and V2 of the 16S rRNA gene were targeted and amplified using Q5 High-fidelity polymerase kit. Primer sequences for amplification are found in **Table S1**. PCR amplification conditions were: 1 cycle of 98°C for 2min, followed by 20 cycles of 98°C for 30s, 50°C for 30s and 72°C for 90s, and a final cycle of 72°C for 5min. PCR products were purified using Ampure XP beads (Agencourt) and pooled in an equivalent molar mix based on the concentration of each PCR product determined by Quant^−^iT™ PicoGreen® dsDNA Assay Kit (Life Tech). Sequencing was performed on an Illumina MiSeq platform using paired-end reads (average read length was 330bp). 16S rRNA gene sequencing analyses and bioinformatics analyses were performed as previously published [25]. All raw reads were deposited in European Nucleotide Archive: PRJEB35291.

### ^1^H NMR analyses

50mg of model colon samples were mixed with 700μL NMR buffer (0.26g NaH2PO, 1.44gK_2_HPO_4_, 17mg TSP, 56.1 mg NaN_3_ and 100 mL D_2_O) and centrifuged at 14000 *xg* at 4°C for 5 min. 600μL of supernatants were transferred into NMR tubes (GPE Scientific Ltd) and the ^1^H NMR spectrum were recorded at 600MHz on a Bruker Avance spectrometer (Bruker BioSpin GmbH, Rheinstetten, Germany) running Topspin 3.2 software. Each ^1^H NMR spectrum was acquired with 128 scans, a spectral width of 12300Hz and an acquisition time of 2.7s. The “noesygppr1d” presaturation sequence was used to suppress the residual water signal with a low^−^power selective irradiation at the water frequency during the recycle delay (3s). Spectra were transformed with a 0.3-Hz line broadening, automatically phased, baseline corrected, and referenced by setting the TSP methyl signal to 0ppm. Metabolites were identified and subsequently quantified using a library of reference standard spectra provided with the Chenomx NMR suite 8.4™ software.

### Statistical analyses and plotting

Statistical analyses on growth curves were performed in GraphPad Prism version 5.04. Mann-Whitney-U-Test was performed for cultures in MRS. Error bars denote standard deviation (SD). All plots were produced using either GraphPad Prism version 5.04 or R Studio version 1.1.463 using the ggplot2 R package version 3.1.0. For NMDS (Non-metric multidimensional scaling) in R Studio the 16S bacterial sequence data was subsampled to an even depth of 120,070 sequences using phyloseq package version 1.24.2. NMDS plots were generated with a Bray^−^Curtis dissimilarity calculation in R Studio used with the vegan package version 2.5^−^4 in R Studio version 1.1463 with R 3.5.0. Spearman correlation analysis was calculated using Kendall in R Studio (version 1.1463).

## Acknowledgements

This work was part funded by an Erasmus studentship to DP. MAEL was funded by the Marie Skłodowska-Curie Individual Fellowship (Project 661594). LJH is funded by a Wellcome Trust Investigator award (100974/C/13/Z) and together with TK by a BBSRC ISP grant for Gut Microbes and Health BB/R012490/1 and its constituent project(s), BBS/E/F/000PR10353 and BBS/E/F/000PR10355. TK is also funded by the Genomics for Food security CSP grant from the BBSRC (BB/CSP17270/1). AT is supported by the BBSRC Norwich Research Park Biosciences Doctoral Training Partnership (grant BB/M011216/1). DvS is supported by Science Foundation Ireland (SFI/12/RC/2273-P1 and SFI/12/RC/2273-P2).

