## Supplementary Figures for "*Bifidobacterium breve* UCC2003 exopolysaccharide modulates the early life microbiota by acting as a dietary substrate"

### Slide 1
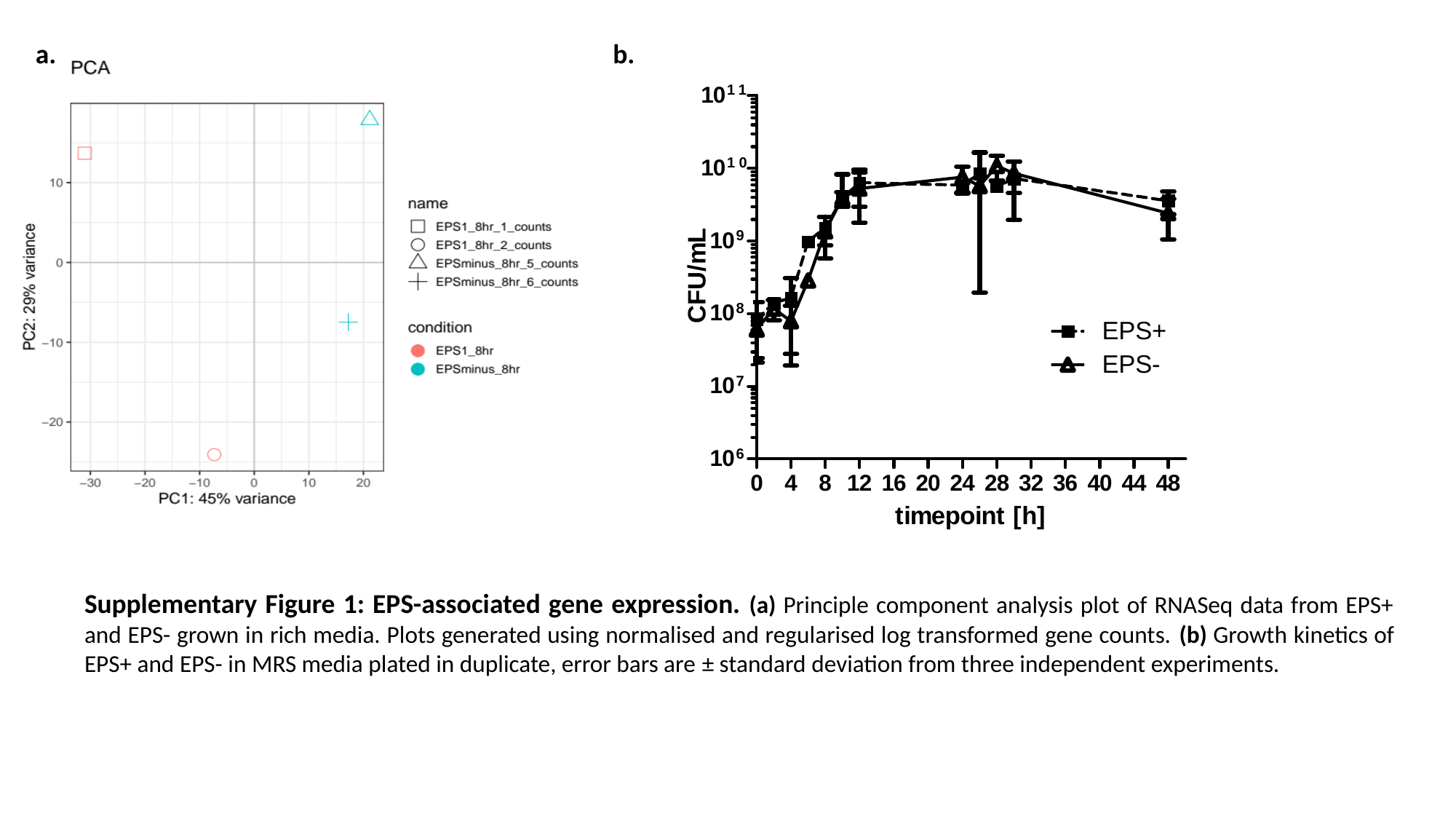

a.
b.
Supplementary Figure 1: EPS-associated gene expression. (a) Principle component analysis plot of RNASeq data from EPS+ and EPS- grown in rich media. Plots generated using normalised and regularised log transformed gene counts. (b) Growth kinetics of EPS+ and EPS- in MRS media plated in duplicate, error bars are ± standard deviation from three independent experiments.

### Slide 2
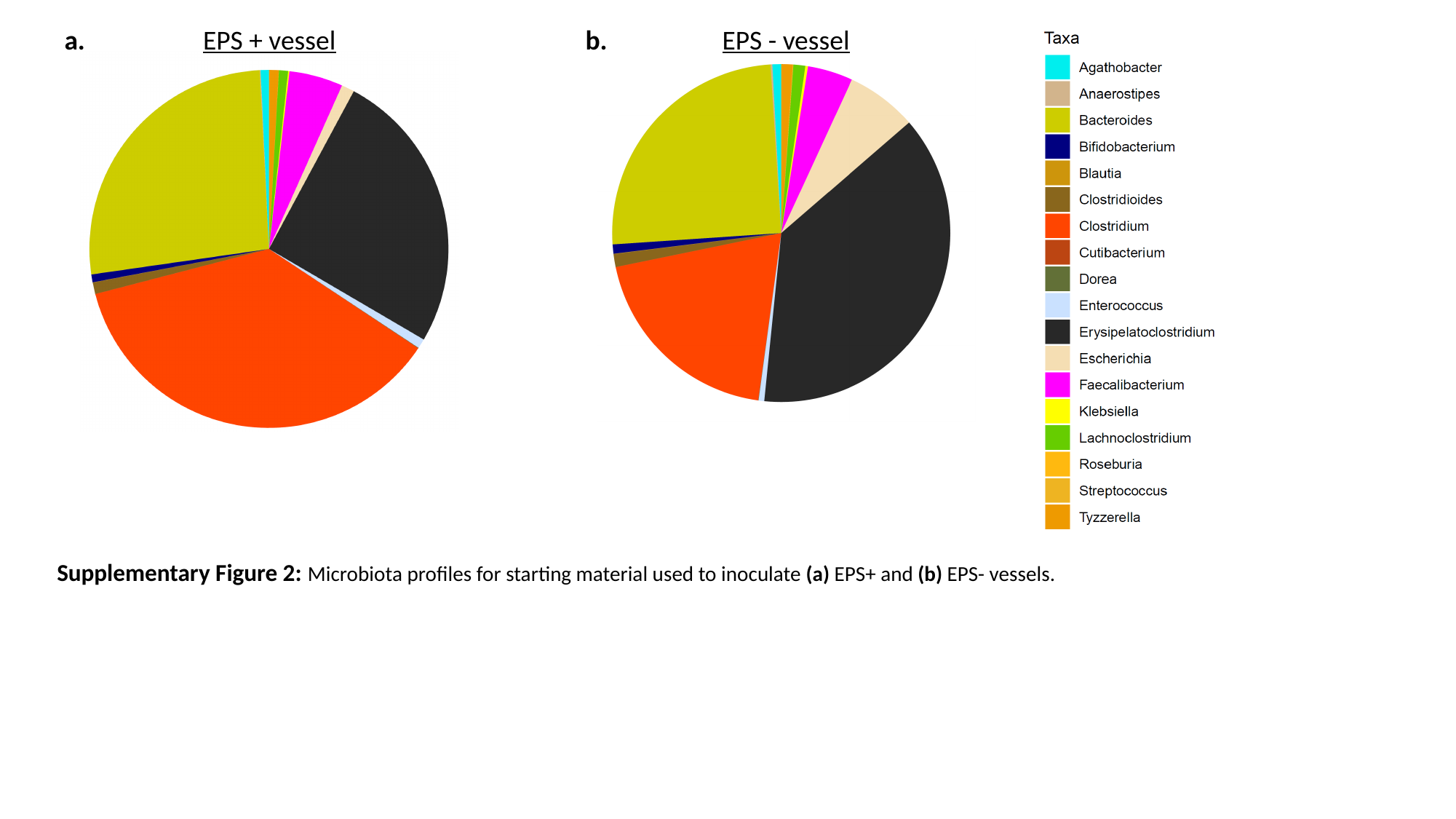

a.
EPS + vessel
b.
EPS - vessel
Supplementary Figure 2: Microbiota profiles for starting material used to inoculate (a) EPS+ and (b) EPS- vessels.

### Slide 3
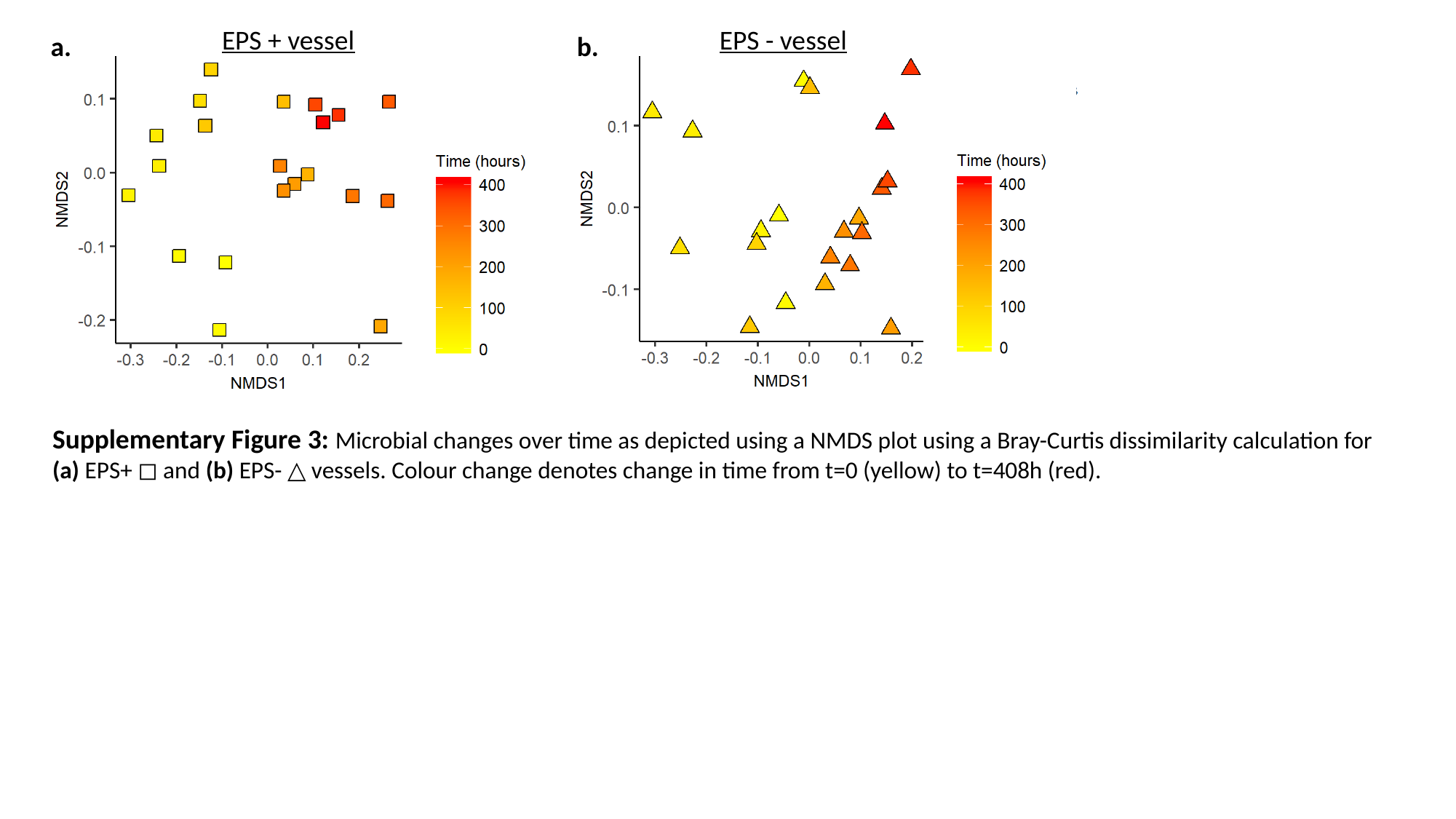

EPS + vessel
EPS - vessel
a.
b.
Supplementary Figure 3: Microbial changes over time as depicted using a NMDS plot using a Bray-Curtis dissimilarity calculation for (a) EPS+ ◻︎ and (b) EPS- △ vessels. Colour change denotes change in time from t=0 (yellow) to t=408h (red).

### Slide 4
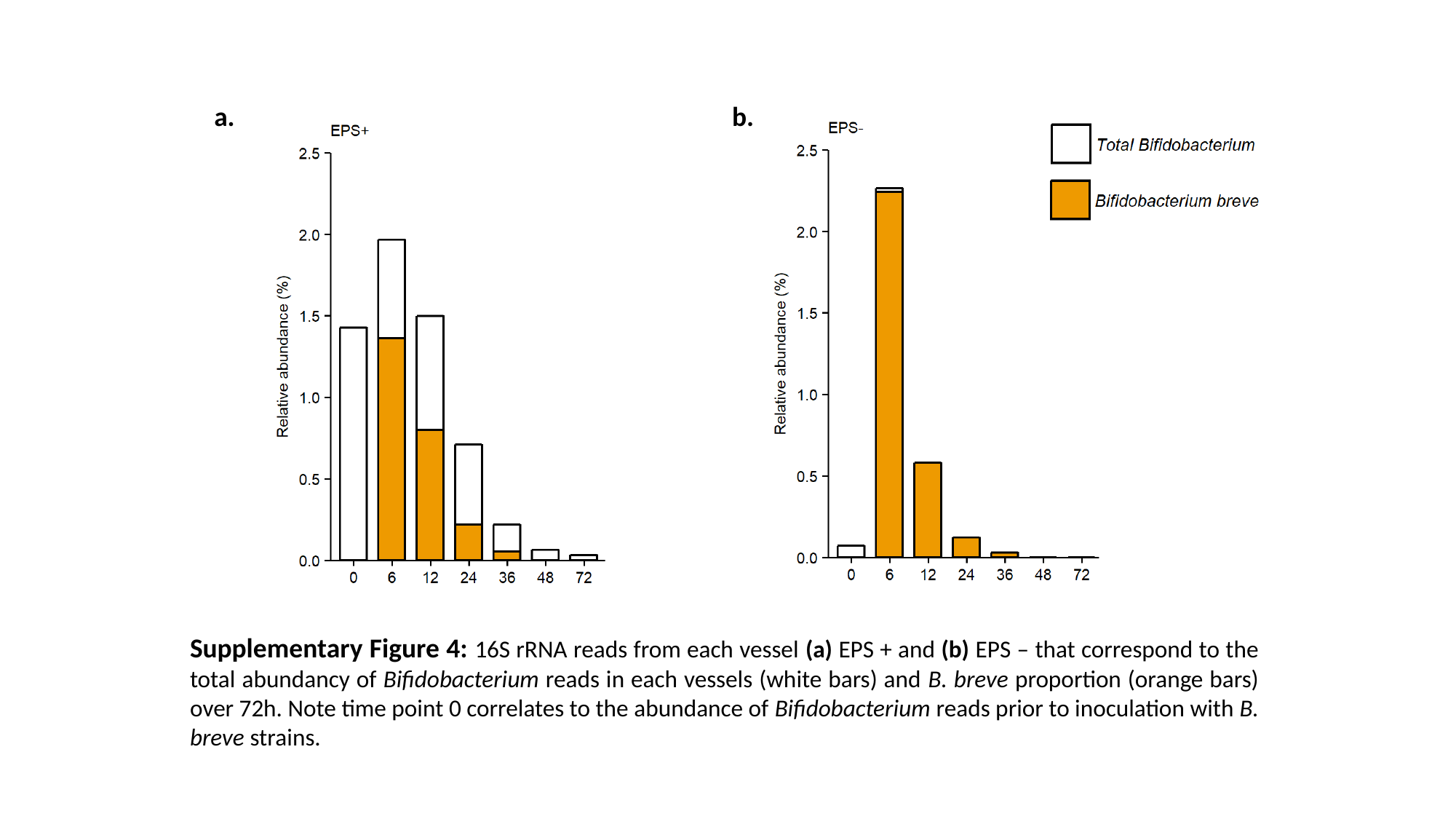

a.
b.
Supplementary Figure 4: 16S rRNA reads from each vessel (a) EPS + and (b) EPS – that correspond to the total abundancy of Bifidobacterium reads in each vessels (white bars) and B. breve proportion (orange bars) over 72h. Note time point 0 correlates to the abundance of Bifidobacterium reads prior to inoculation with B. breve strains.

### Slide 5
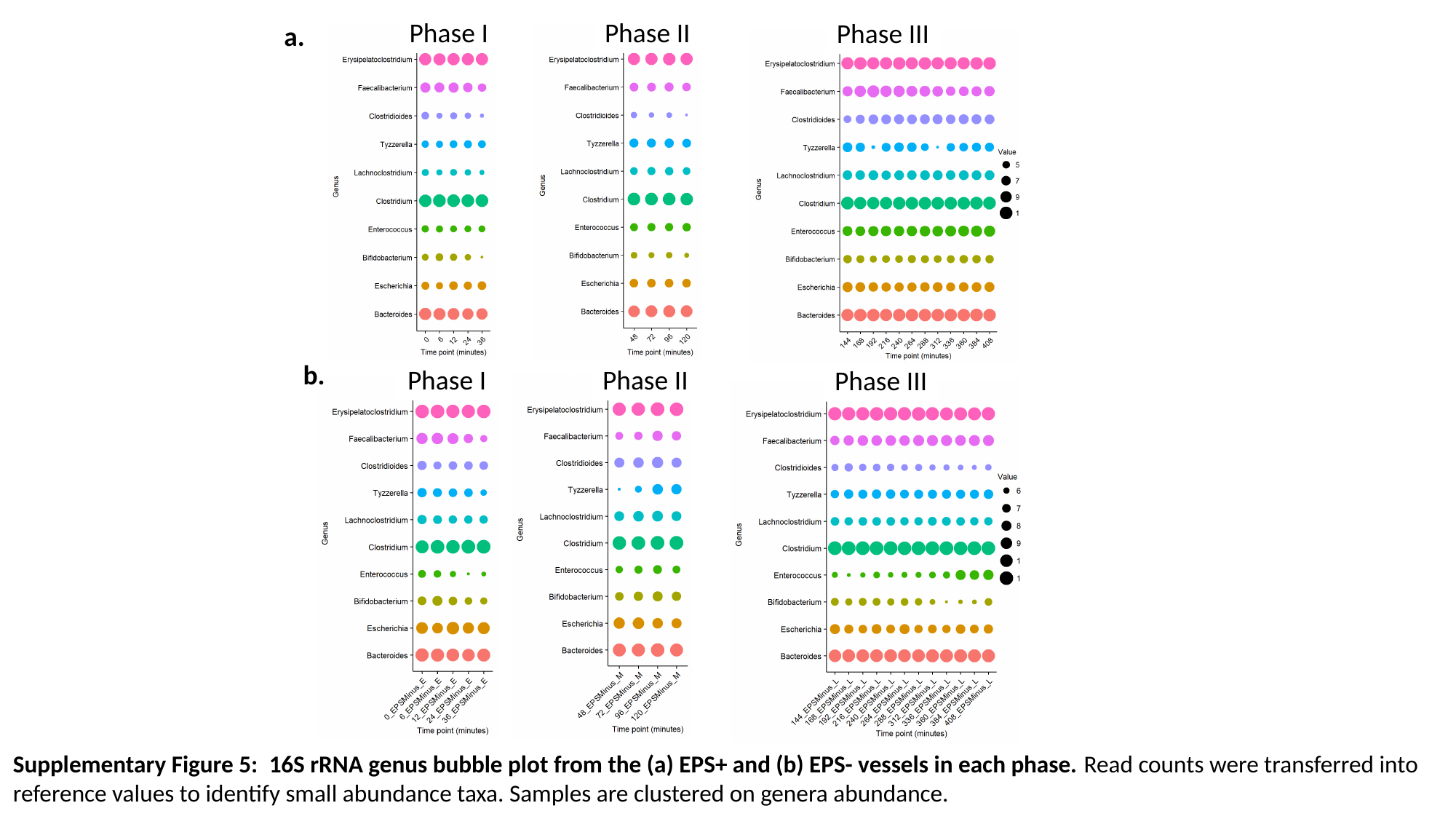

Phase I
Phase II
Phase III
a.
b.
Phase I
Phase II
Phase III
Supplementary Figure 5: 16S rRNA genus bubble plot from the (a) EPS+ and (b) EPS- vessels in each phase. Read counts were transferred into reference values to identify small abundance taxa. Samples are clustered on genera abundance.

### Slide 6
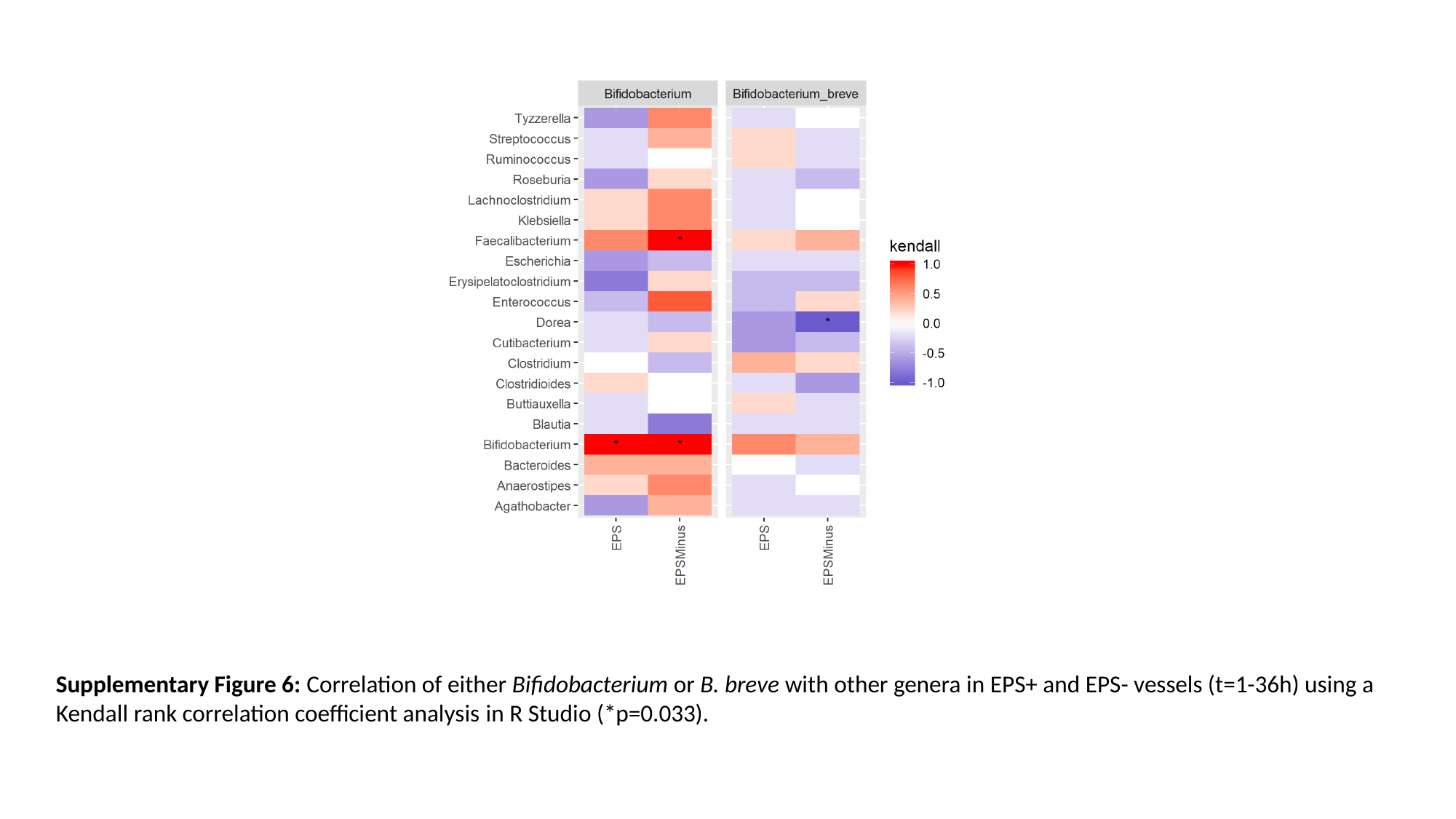

Supplementary Figure 6: Correlation of either Bifidobacterium or B. breve with other genera in EPS+ and EPS- vessels (t=1-36h) using a Kendall rank correlation coefficient analysis in R Studio (*p=0.033).

### Slide 7
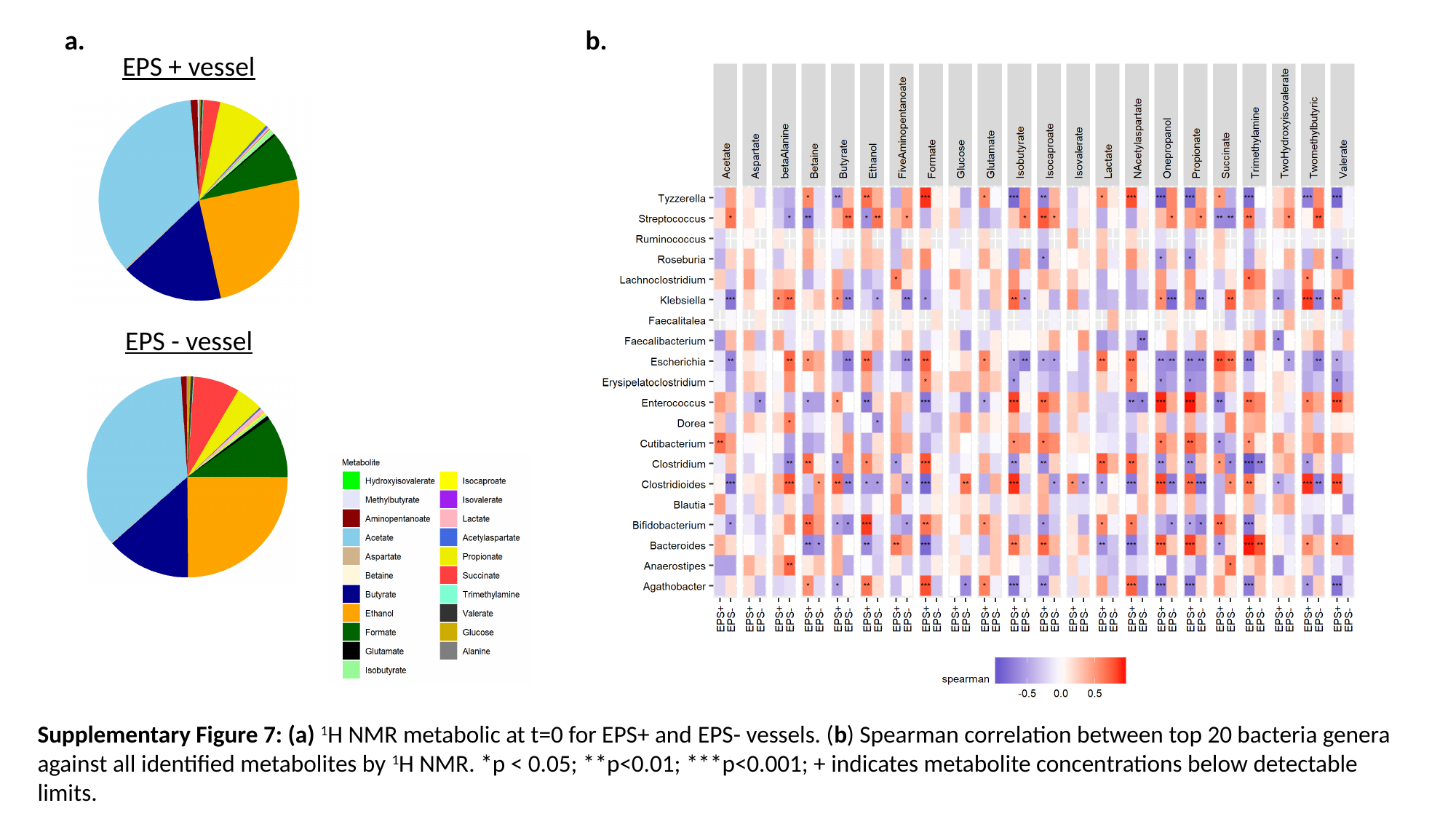

a.
b.
EPS + vessel
EPS - vessel
Supplementary Figure 7: (a) ^1^H NMR metabolic at t=0 for EPS+ and EPS- vessels. (b) Spearman correlation between top 20 bacteria genera against all identified metabolites by ^1^H NMR. *p < 0.05; **p<0.01; ***p<0.001; + indicates metabolite concentrations below detectable limits.

### Slide 8
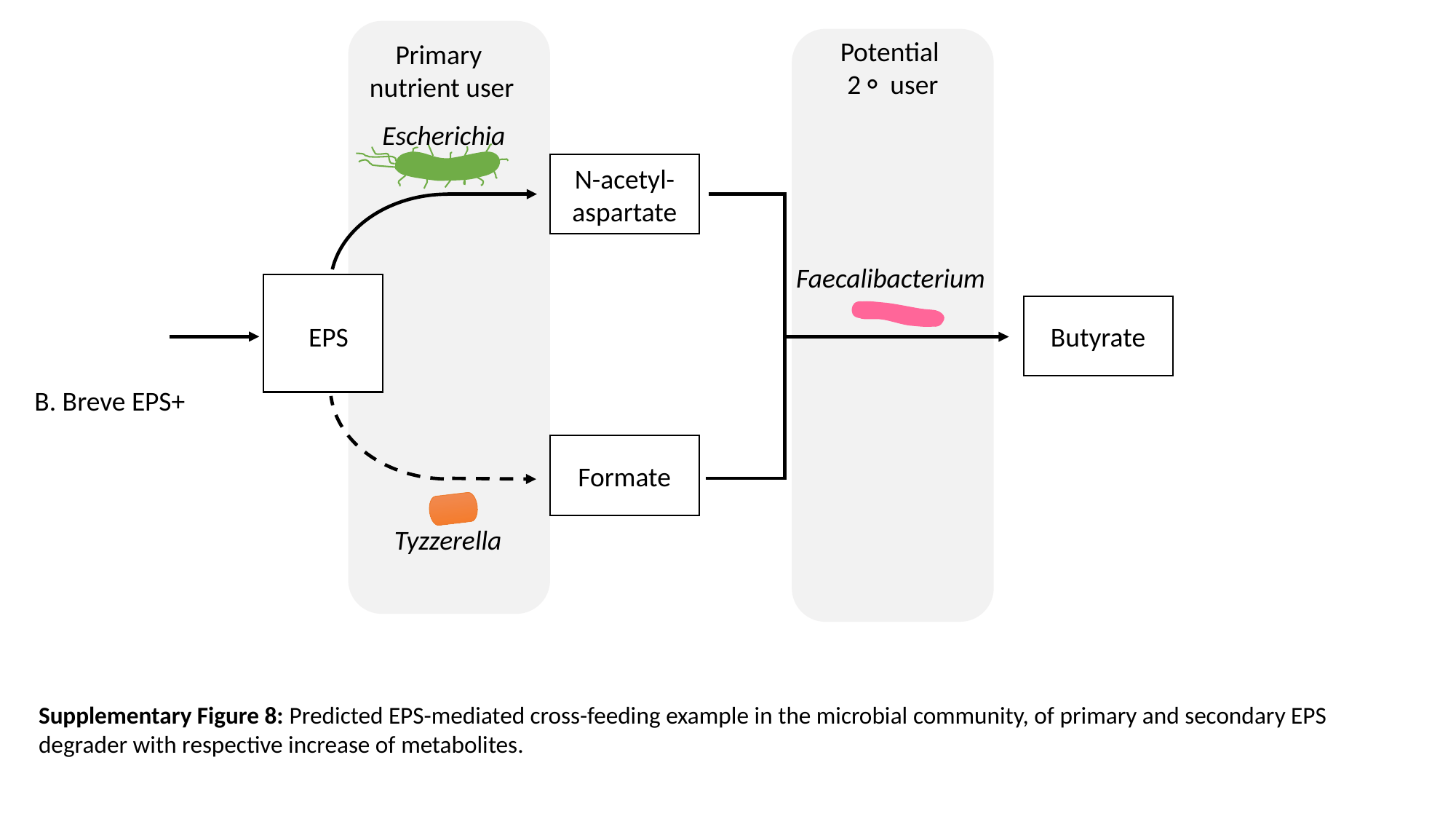

Potential
2⚬ user
Primary
nutrient user
Escherichia
N-acetyl-aspartate
Faecalibacterium
Butyrate
EPS
B. Breve EPS+
Formate
Tyzzerella
Supplementary Figure 8: Predicted EPS-mediated cross-feeding example in the microbial community, of primary and secondary EPS degrader with respective increase of metabolites.
